# Heart Slice Culture System Reliably Demonstrates Clinical Drug-Related Cardiotoxicity

**DOI:** 10.1101/2020.06.12.148197

**Authors:** Jessica M. Miller, Moustafa H. Meki, Qinghui Ou, Sharon A. George, Anna Gams, Riham R. E. Abouleisa, Xian-Liang Tang, Brooke M. Ahern, Guruprasad A. Giridharan, Ayman El-Baz, Bradford G. Hill, Jonathan Satin, Daniel J. Conklin, Javid Moslehi, Roberto Bolli, Alexandre J. S. Ribeiro, Igor R. Efimov, Tamer M. A. Mohamed

**Author notes:** Correspondence to: Tamer M A Mohamed, Institute of Molecular Cardiology, University of Louisville, 580 South Preston Street, Louisville, KY 40202., Igor R Efimov, Department of Biomedical Engineering, The George Washington University, 5000 Science and Engineering Hall, Washington, DC 20052, USA., Alexandre J. S. Ribeiro, U.S. Food and Drug Administration, Center for Drug Evaluation and Research, Office of Translational Science, Office of Clinical Pharmacology, Division of Applied Regulatory Science, Silver Spring, MD, USA. Authors equally contributed to the work (Jessica M. Miller, Moustafa H. Meki, Qinghui Ou, Sharon A. George).

## Abstract

The limited availability of human heart tissue and its complex cell composition are major limiting factors for reliable testing drug efficacy, toxicity and understanding mechanism. Recently, we developed a functional human and pig heart slice biomimetic culture system that fully preserves the viability and functionality of 300 µm heart slices for 6 days. Here, we tested the reliability of this culture system in delineating the mechanisms of known anti-cancer drugs that cause cardiomyopathy. We tested three anti-cancer drugs (doxorubicin, trastuzumab, and sunitinib) associated with different mechanisms leading to cardiotoxicity at three concentrations and assessed the effect of these drugs on heart slice viability, structure, function and transcriptome. Slices incubated with any of these drugs for 48 h showed significant loss in viability, cardiomyocyte structure and functionality. Mechanistically, RNA sequencing demonstrated a significant downregulation of cardiac genes and upregulation of oxidative response in doxorubicin-treated tissues. Trastuzumab treatment caused major downregulation in cardiac muscle contraction-related genes, consistent with its clinically known direct effect on cardiomyocytes. Interestingly, sunitinib treatment resulted in significant downregulation of angiogenesis-related genes in line with its mechanism of action. Heart slices are not only able to demonstrate the expected toxicity of doxorubicin and trastuzumab similar to hiPS-derived-cardiomyocytes; they are superior in detecting sunitinib cardiotoxicity phenotypes and mechanism in the clinically relevant concentration range, 100 nM – 1 µM. These results indicate that heart slice tissue culture models have the potential to become a reliable platform for testing drug toxicity and mechanism of action.

## Introduction

In the past 30 years, cardiotoxicity induced by side effects of drugs that affect the electrical activity as well as the contractility of cardiomyocytes has been the cause for 40% of all drug withdrawals from the market^1, 2^. In addition, in the last decade, there has been an explosion of cancer therapies, several of which often lead to cardiotoxic effects, causing cardiomyopathy, arrhythmias, irreversible heart failure or death^3^. For example, both traditional (e.g., anthracyclines and radiation) and targeted (e.g., trastuzumab) breast cancer therapies can result in cardiovascular complications in a subset of patients^4, 5^. The cardiovascular effects of newer classes of drugs, such as CDK4/6 inhibitors and PI3K inhibitors, are still unclear^6, 7^. The recent higher cancer survival is partially offset by increased morbidity and mortality related to the cardiotoxic side effects of anti-cancer therapeutics^8^. Hence, a close collaboration between cardiologists and oncologists (in the emerging field of cardio-oncology) aims to make these complications manageable to ensure that patients can be treated as safely and as effectively as possible^2^.

Detection of cardiotoxic effects of drug candidates requires the use of *in vivo* and *in vitro* studies prior to clinical trials^9^. Therefore, there is a growing need for reliable preclinical screening strategies for cardiovascular toxicities associated with emerging cancer therapies prior to human clinical trials. Animal models can fail in predicting drug adverse cardiac effects^10, 11^, can be expensive and do not replicate many of the biochemical properties and hemodynamic aspects of the human heart and circulation^12-14^. Recently, human induced pluripotent stem cell derived cardiomyocytes (hiPSC-CMs) started being used to assess cardiotoxicity^15, 16^. However, these cells have several fetal-like properties that can limit their usefulness in comprehensively predicting clinical cardiac drug side effects. The development of hiPSC-CMs microtissues is a promising effort to generate more mature phenotypes, but it is still a work in progress^15, 16^; because such microtissues still exhibit immature electrophysiology and lack cell-cell electrical coupling^17^. A less common approach has been the use of isolated primary human cardiomyocytes. While these cells are functionally mature, and can be used for high-throughput testing, they readily dedifferentiate in culture, thereby limiting their use for cardiotoxicity studies^18^. The adult human heart tissue is structurally more complicated, being composed of a heterogeneous mixture of various cell types including cardiomyocytes, endothelial cells, smooth muscle cells, and various types of stromal fibroblasts linked together with a sophisticated three-dimensional network of extracellular matrix proteins^19^. This heterogeneity of the non-cardiomyocyte cell population^20-22^ in the adult heart is a major obstacle in modeling heart tissue using individual cell types. Human ventricular wedge preparations have also been used to study electrophysiology^23, 24^. However, the wedge preparations are typically large in size and have functional viability of only several hours, limiting their applicability to high-throughput applications.

These major limitations highlight the importance of developing optimal methods to enable the culture of intact cardiac tissue for studies involving physiological and pathological conditions^17^. Culturing human heart slices has shown promise in extending the functionality and viability of cardiac tissue. By retaining the normal tissue architecture, the multi-cell type environment, and the 3-dimensional structure, tissue slices can more faithfully replicate the organ-level adult cardiac physiology in terms of normal conduction velocity, action potential duration, and intracellular calcium dynamics^25, 26^. Recently, we have developed a novel biomimetic culture system that maintains full viability and functionality of human and pig heart slices (300 µm thickness) for 6 days in culture through optimization of the medium and culture conditions with continuous electrical stimulation at 1.2 Hz and oxygenation of the culture medium^27^. In the present study, we aim to test the reliability of this culture system in recapitulating clinical toxicity and potentially delineating the mechanisms of known cardiotoxic drugs. To this end, we selected three anticancer drugs with known adverse cardiac side effects: doxorubicin, an anthracycline with a long history of cardiotoxicity; trastuzumab, a monoclonal antibody target HER2; and, sunitinib, a small molecule multi-targeted kinase inhibitor (TKI) with potent activity against VEGF and PDGF receptors^5^. All three drugs have been associated with cardiomyopathy through a different mechanisms^5^. These 3 drugs were selected as in hiPSC-CMs doxorubicin and trastuzumab are shown to be cardiotoxic^28, 29^, however, nanomolar concentrations of sunitinib do not cause any cardiotoxicity on hiPSC-CMs ^30^. Therefore, we first confirmed in our hands the previously reported data that hiPSC-CMs are able to demonstrate the cardiotoxic phenotype of doxorubicin, but not sunitinib, at low concentrations^30^. Then, in pig heart slices, we tested three different concentrations of each drug in ten-fold incremental values with the lowest dose being the most clinically relevant and assessed the effect of these drugs on heart slice viability, structure, function and transcriptome. As the maximum blood concentration of doxorubicin ranges between 100nM-300nM^31^, we used 100nM, 1µM, 10µM concentrations.

We used 1µg/mL, 10µg/mL and 100µg/mL concentrations for trastuzumab as the blood level is ranging between 1-10µg/mL^32^. Sunitinib concentrations of 100nM, 1µM, 10µM were used as the normal blood concentration range is 100nM-200nM^33^. Finally, we assessed the ability and reproducibility of the human heart slices in demonstrating the true cardiotoxic mechanism of doxorubicin between different human subjects.

## Methods

### Apoptotic assay on hiPSC-CMs

hiPSC-CMs (iCell Cardiomyocytes^2^) were purchased from Cellular Dynamics (Currently FujiFilm Inc.). These hiPSC-CMs have been selected after differentiation using an a-MHC-Blastocidin selection cassette. This strategy yields nearly 100% pure hiPSC-CMs. Following vendor recommendations, hiPSC-CMs were seeded on fibronectin coated plates and cultured in maintenance medium.. hiPSC-CMs were exposed to a range of concentrations of doxorubicin (cardiotoxin), sunitinib (cardiotoxic TKI), aspirin (non-toxic drug), erlotinib (non-cardiotoxic TKI) over 48 h. DMSO in the medium at a concentration of 0.1% (v/v) was used as a control condition for each drug. Live cell brightfield and fluorescent images for time-lapse experiments were acquired using the Incucyte® S3 Live Cell Analysis System (Essen Bioscience / Sartorius) running version 2019A (20191.1.6976.19779). Plates were imaged hourly following initiation of drug treatment. Three images at set locations within each well were acquired at 10x magnification for each time point. For fluorescent imaging of apoptosis, the Incucyte Caspase-3/7 Green dye (Essen Bioscience 4440) was added to the media/drug cocktail at a concentration of 5 μM for the duration of the assay. Analysis of apoptosis using the Caspase 3/7 fluorescent dye was performed using the Incucyte software. Parameters were set to quantify green fluorescent intensity metrics and areas of high signal intensity are marking cell death.

### Harvesting porcine heart tissue

All animal procedures were in accordance with the institutional guidelines and approved by the University of Louisville Institutional Animal Care and Use Committee. The protocol for harvesting pig hearts has been described in detail^27, 34^. Briefly, following deeply anesthetizing the pig with 5% isoflurane, pig heart was quickly excised out and the heart was clamped at the aortic arch and perfused with 1L sterile cardioplegia solution (110 mM NaCl, 1.2 mM CaCl_2_, 16 mM KCl, 16 mM MgCl_2_, 10 mM NaHCO_3_, 5 units/mL heparin, pH to 7.4), then the heart was preserved on an ice-cold cardioplegic solution and immediately transported to the lab on wet ice.

### Human heart Slices

Donor hearts that were not used in transplantation were acquired through our collaboration with the Washington Regional Transplant Community. Human cardiac organotypic slices were prepared as previously described^35, 36^.

### Heart slicing and culturing

Slicing and culturing 300 µm thick heart tissue slices were performed as previously described in^27, 34^. A refined oxygenated growth medium was used (Medium 199, 1x ITS supplement, 10% FBS, 5 ng/mL VEGF, 10 ng/mL FGF-basic, and 2x Antibiotic-Antimycotic), and changed 3 times/day. Sunitinib (Tocris Inc.) (100 nM, 1µM, and 10 µM), trastuzumab (InvivoGen Inc.) (1 µg, 10 µg, and 100 µg), or doxorubicin (Sigma Millipore Inc.) (100 nM, 1µM, and 10 µM, and 50 µM) was added freshly to the culture medium at each medium change. Control slices received DMSO at the same dilution factor as the drug-treated slices.

### MTT viability assay

For the MTT assay we used the Vybrant ® MTT Cell Proliferation Assay kit (Thermo Scientific) following the manufacturer’s protocol with slight modifications. Using a sterile scalpel, each heart slice was cut into 2 ∼0.5 cm^2^-sized sections for the MTT assay. The heart slice segments were placed into a single well of a 12-well plate containing 0.9 mL growth media with 0.1 mL of reconstituted MTT substrate that was prepared according to the manufacturer’s protocol. The tissue was incubated at 37°C for 3 h. During this time, viable tissue metabolized the MTT substrate and produced a purple color formazan compound. To extract the purple formazan from the tissue slices, the tissue was transferred into 1 mL of DMSO and incubated at 37°C for 15 min. The resulting solution was purple in color. This solution was then transferred into a clear bottom 96-well plate in triplicate at 3 dilutions, 1:2, 1:5, and 1:10. The intensity of the purple color was measured using cytation 1 plate reader (BioTek) at 570 nm. The readings were normalized to the weight of each heart tissue section and converted into OD/mg tissue. The 3 dilutions of the solution were performed to correct for any possible signal saturation and the average of all readings was normalized to their dilution factor.

### Heart slice fixation, mounting and immunofluorescence

Heart slices were fixed with 4% paraformaldehyde for 48 h. Fixed tissue was dehydrated in 10% sucrose for 1 hour, 20% sucrose for 1 h, and 30% for overnight. The dehydrated tissue was then embedded in optimal cutting temperature compound (OCT compound) and gradually frozen in isopentane/dry ice bath. OCT embedded blocks were stored at -80°C until sectioning. 8 µm sections were cut and immunolabeled for target proteins using the following procedure: To remove the OCT compound, the slides were heated for 5 min at 95°C until the OCT compound melted. Then 1 mL of PBS was added to each slide and incubated at RT for 10-30 min until the OCT compound washed off. Sections were then permeabilized by incubating for 30 min in 0.1% Triton-X in PBS at RT. The Triton-X was removed, and non-specific antibody binding of the sections were blocked with 3% BSA solution for 1 h at RT. After washing the BSA off with PBS, each section was marked off with a wax pen. After marking sections off, the primary antibodies (1:200 dilution in 1% BSA) connexin 43 (Abcam; #AB11370), troponin-T (Thermo Scientific; #MA5-12960)] were added to each section and incubated for 90 min at RT. The primary antibodies were washed off with PBS three times followed by the addition of the secondary antibodies (1:200 dilution in 1% BSA) anti-Ms AlexaFluor 488 (Thermo Scientific; #A16079), anti-Rb AlexaFluor 594 (Thermo Scientific; #T6391) and incubated for 90 min at RT. The secondary antibody was removed by washing the sections 3 times with PBS. To distinguish the *bona fide* target staining from the background, we used a secondary antibody only as a control. After 3 times PBS washes, DAPI was added for 15 min. The sections were washed again 3 times with PBS. Finally, slices were mounted in vectashield (Vector Laboratories) and sealed with nail polish. All immunofluorescence imaging and quantification were performed using a Cytation 1 high content imager and the fluorescent signal quantification and masking were performed using the Gen5 software.

### Calcium-transient assessment

Calcium transients were assessed as previously described^27^. Briefly, heart slices were loaded with Fluo-4 for 30 min at room temperature before being transferred to the imaging chamber. The loading solution contained a 1:10 mixture of 5 mM Fluo-4 AM in dry DMSO and PowerloadTM concentrate (Invitrogen), which was diluted 100-fold into extracellular Tyrode’s solution (NaCl 140mM; KCl 4.5mM; glucose 10mM; HEPES 10mM; MgCl_2_ 1mM; CaCl_2_ 1.8mM; 2x Antibiotic-Antimycotic; pH 7.4). Additional 20 min were allowed for de-esterification before recordings were taken. Contractions and calcium transients were evoked by applying voltage pulses at 1 Hz between platinum wires placed on either side of the heart slice and connected to a field stimulator (IonOptix, Myopacer). Fluo-4 fluorescence transients were recorded via a standard filter set (#49011 ET, Chroma Technology). Resting fluorescence was recorded after cessation of pacing, and background light was obtained after removing the heart slice from the field of view at the end of the experiment. All analyses of calcium transients were based on calcium transients recorded from single cardiomyocytes within the heart slice and the calcium transient’s amplitude was assessed as the average of 10 consecutive beats from each cell. Calcium transients and amplitude was assessed following normalization to the basal florescence of each cell and represented as F/F_0_.

### RNA Sequencing

RNA was isolated from the heart slices by using the Qiagen miRNeasy Micro Kit, #210874, following the manufacturer’s protocol after homogenization of tissue in Trizol. RNAseq library preparation, sequencing, and data analysis were performed as described previously^27^.

### Optical Mapping

Slices were incubated with Di-4-ANEPPS (30 µl of stock solution at 1.25 mg/mL diluted to 1 mL with recovery solution(140 mM NaCl, 4.5 mM KCl, 1 mM MgCl2, 1.8 mM CaCl2, 10 mM glucose, 10 mM HEPES, 10 mM BDM; pH 7.4)) for single parameter voltage optical mapping. Slices were paced at 1 Hz frequency, 2 ms duration at an amplitude of 1.5X the threshold for stimulation using a platinum bipolar pacing wire positioned at the center of the slice. Slices were excited using excitation light at 520 ± 5 nm and the emitted light was collected by a tandem lens optical mapping system, filtered using a 610 ± 20 nm filter and recorded using the SciMedia Ultima L-type CMOS cameras.

Optical signals were analyzed as previously described^35, 36^. Briefly, activation times were defined as the time of maximum first derivative of the fluorescence signal during the upstroke. End of action potential and calcium transients were defined as the point where the signal returns to 80% of its total signal amplitude. Conduction velocity was determined in the transverse direction of propagation using activation times and known interpixel resolution.

### Cap analysis of gene expression

For high-throughput analysis of transcriptional starting point and identification of promoter usage, we performed cap analysis of gene expression (CAGE). Total RNA from the cultured slices was extracted using the RNeasy Fibrous Tissue Mini Kit according to the manufacture’s protocol. Library preparation and sequencing with HiSeq2500 was performed according to the Morioka *et al* protocol^37^. Read mapping was performed with Burrows-Wheeler Aligner^38^; subsequent analysis of reads into decomposition peak identification (DPI) and transcription start site-like peaks was performed with *DPI1* (https://github.com/hkawaji/dpi1/) followed by *TomeTools TSSClassifier* (https://sourceforge.net/projects/tometools/). Differential expression analysis was performed with *edgeR*^*39*^, gene ontology (GO) with *limma*^40^, and heatmap plotting with *pheatmap* (https://CRAN.R-project.org/package=pheatmap).

## Results

### hiPSC-CMs demonstrate doxorubicin cardiotoxicity but not sunitinib

A major distinction between the heart slice preparation and hiPSC-CMs is the retention of multiple cell types in heart slices. It has been reported hiPSC-CMs is able to demonstrate the cardiotoxic phenotype of doxorubicin in nanomolar concentrations but not for nanomolar concentrations of sunitinib^30^. First we needed to reproduce these findings in our hands, therefore, we tested different the effect of different concentrations of doxorubicin, and sunitinib on hiPS-CMs viability using caspapse3/7 apoptosis assay over 48 hours. aspirin and erlotinib (non-cardiotoxic TKI) used as negative controls (Figure 1). consistent with the previous report^30^, sunitinib did not show any cardiotoxicity in hiPSC-CMs at 500 nM or 1 µM concentrations as assessed by caspase3/7 apoptosis assay (Figure 1d-f). Sunitinib at 60 µM showed acute apoptotic response within one hour of addition, therefore, no more fluorescence was detected over the course of the experiment using this concentration (Figure 1d-f). However, doxorubicin as low as 500 nM demonstrated the expected cardiotoxicity on hiPSC-CMs and the negative controls (aspirin and erlotinib (non-cardiotoxic TKI) did not show any cardiotoxic effect on hiPSC-CMs up to 60 µM (Figure 1a-c).

**Figure 1.**
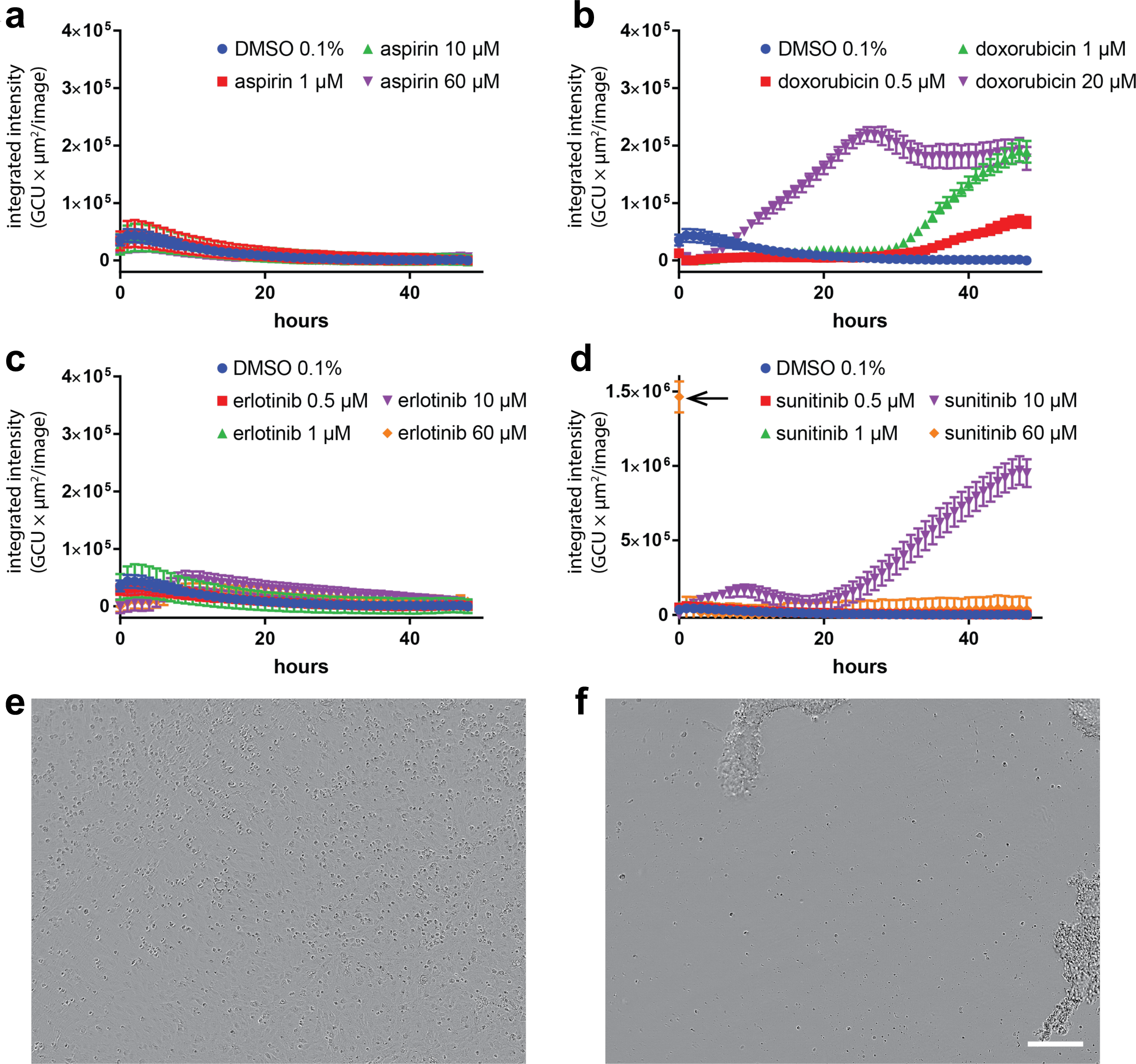
Pro-apoptotic effect of doxorubicin and sunitinib on hiPSC-CMs: Kinetics of caspase-3/7 activation in hiPSC-CMs exposed to a range of concentrations of **(a)** aspirin (non-toxic drug), **(b)** doxorubicin (cardiotoxin), **(c)** erlotinib (non-cardiotoxic TKI) and **(d)** sunitinib (cardiotoxic TKI) over 48 h. DMSO in the medium at a concentration of 0.1% (v/v) was used as a control. (A-D) Variations in total caspase integrated intensity was calculated from fluorescent images of the caspase-3/7 green dye acquired every hour, which captured the extent of apoptosis within cultured cells. Acute apoptotic effects were detected upon exposing cells to 60 µM of sunitinib as noted by the arrow in **(d). (e and f)** Phase microscopy images of cells acquired when 60 µM of sunitinib was added to cells **(e)** and one hour after being exposure **(f)**, where total cell detachment was observed. Scale bar: 200 µm.

### Effect of cardiotoxins on heart slices viability and cardiomyocyte integrity

Pig heart slice exposure to these concentrations of cardiotoxins for 48 h resulted in a significant decline in the general viability of the tissue as assessed by the MTT viability assay (Figure 2a). Structurally, following the tissue exposure to doxorubicin, there was a decrease in the gap junction protein, connexin-43, expression starting at the lowest concentration (Figure 1b). Heart slices treated with trastuzumab showed disruption in connexin-43 localization at the gap junction accompanied by a lower level of troponin expression starting at the lowest concentration (Figure 2b). Interestingly, low concentration sunitinib treatment did not appear to have an effect on the structure of the tissue slices; however, the higher concentrations showed disruption of connexin-43 localization (Figure 2b).

**Figure 2.**
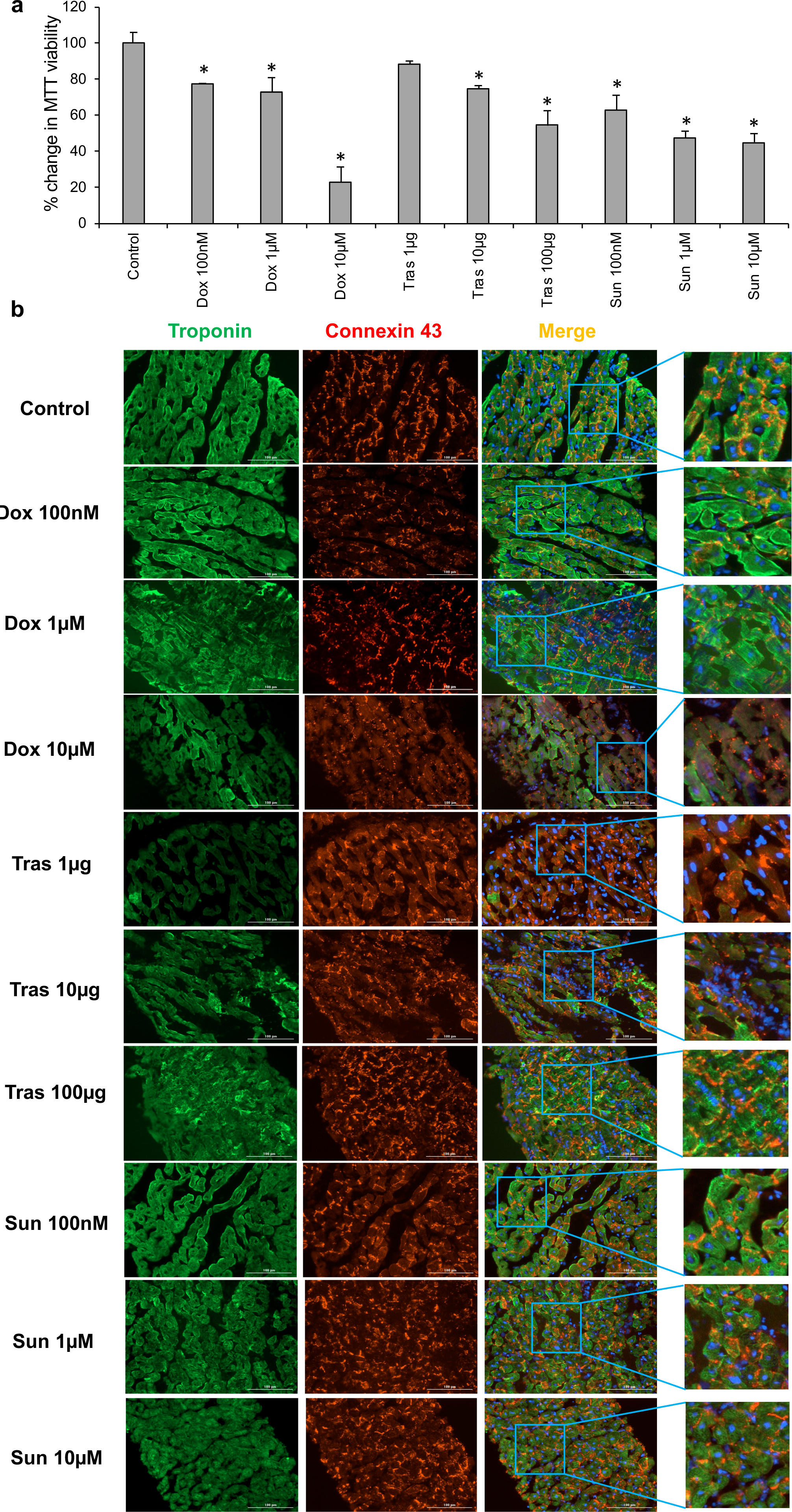
Effect of cardiotoxins on heart slice viability and structure: **(a)** Bar graph shows the quantification of heart slice viability after 2 days of treatment with the corresponding cardiotoxin using MTT assay (n=2 independent experiments, 4 replicates in each, One-Way ANOVA test was conducted to compare between groups; *P<0.05 compared to the control). **(b)** Representative immunofluorescence images showing the expression of connexin 43 (red) in cardiomyocytes (green) in cross sections taken from heart tissue slices treated for 2 days with the corresponding concentration of the cardiotoxin (Scale bar, 100 µm). These representative images have been reproducible over 2 independent experiments with 3 technical replicates in each experiment; however, the lack of reliable tools for quantifying localization of connexin 43 has limited our ability for quantification.

### Effect of cardiotoxins on heart slices calcium homeostasis

We next tested the effects of cardiotoxins on the calcium homeostasis within the pig heart slices after 48 h exposure. All high concentrations of cardiotoxins completely abolished the calcium transients (Supp Movie 3, 4, 6, 7, 9 and 10) compared with the control (Supp movie 1), However, as the lower concentrations are more clinically relevant, they demonstrated several of the clinically-observed effects on the cardiac contractility and rhythm^5^. Treatment with 100 nM doxorubicin showed abolished calcium transients, indicating direct cardiomyocyte damage (Figure 3 Supp movie 2). Trastuzumab (1 µg) (Figure 3, Supp movie 5) or sunitinib (100 nM) (Figure 3, Supp movie 8) treatment resulted in severe disruption in calcium homeostasis within the heart slices.

**Figure 3.**
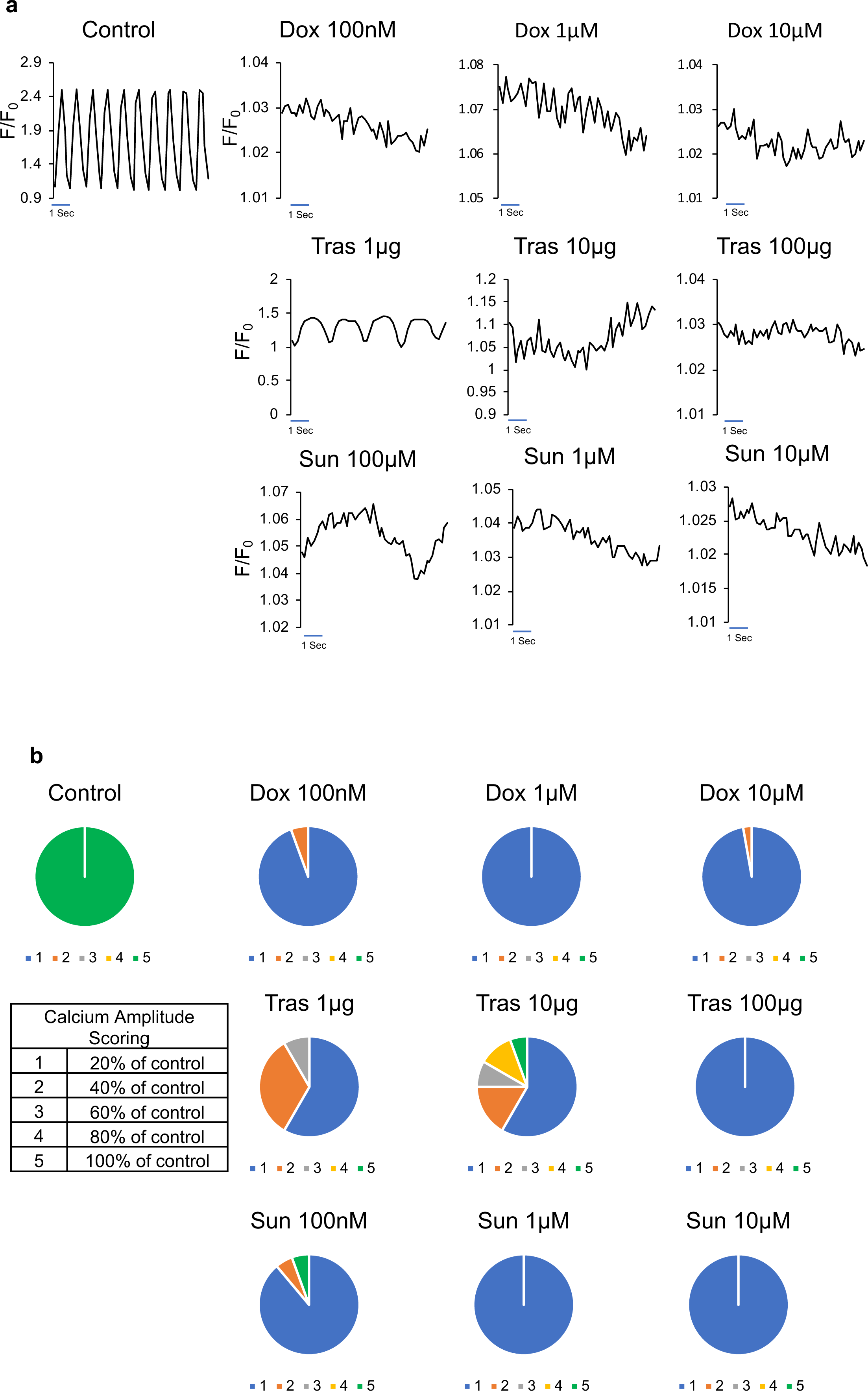
Effect of cardiotoxins on heart slices functionality and calcium homeostasis: **(a)** Representative calcium traces from day 2 cultured control slices and heart slices treated with the corresponding concentration of each cardiotoxin for 2 days. Transients were recorded after loading the heart slices with Fluo-4 calcium dye and using 1 Hz/20 V electrical stimulation at the time of recording. **(b)** Scoring of calcium transient amplitude as indication of the cardiomyocyte function from slices treated with each cardiotoxin (n=36 cells in each group from 2 independent experiments).

### Mechanistic understanding of the cardiotoxic effects

Transcriptomic analyses were performed to understand the mechanism behind the responses of the pig heart slice to cardiotoxins. The top significant up/downregulated gene ontology (GO) terms for tissue treated with 100 nM of doxorubicin are shown in Figure 4a-c. The top downregulated GO terms were genes responsible for cardiac muscle and development as well as cellular division genes. The top upregulated GO terms were genes involved in oxidation/reduction and inflammatory response that is consistent with the known free radical induction by doxorubicin. Figure 5a-c shows the up/downregulated genes in heart slices treated with 1 µg trastuzumab for 48 h. The downregulated genes are mainly contractile genes, which indicates a direct effect on the cardiomyocytes structure. Figure 6a-c shows the up/downregulated GO terms in heart slices treated with 100 nM sunitinib. Sunitinib treatment resulted in significant downregulation in angiogenesis-related genes.

**Figure 4.**
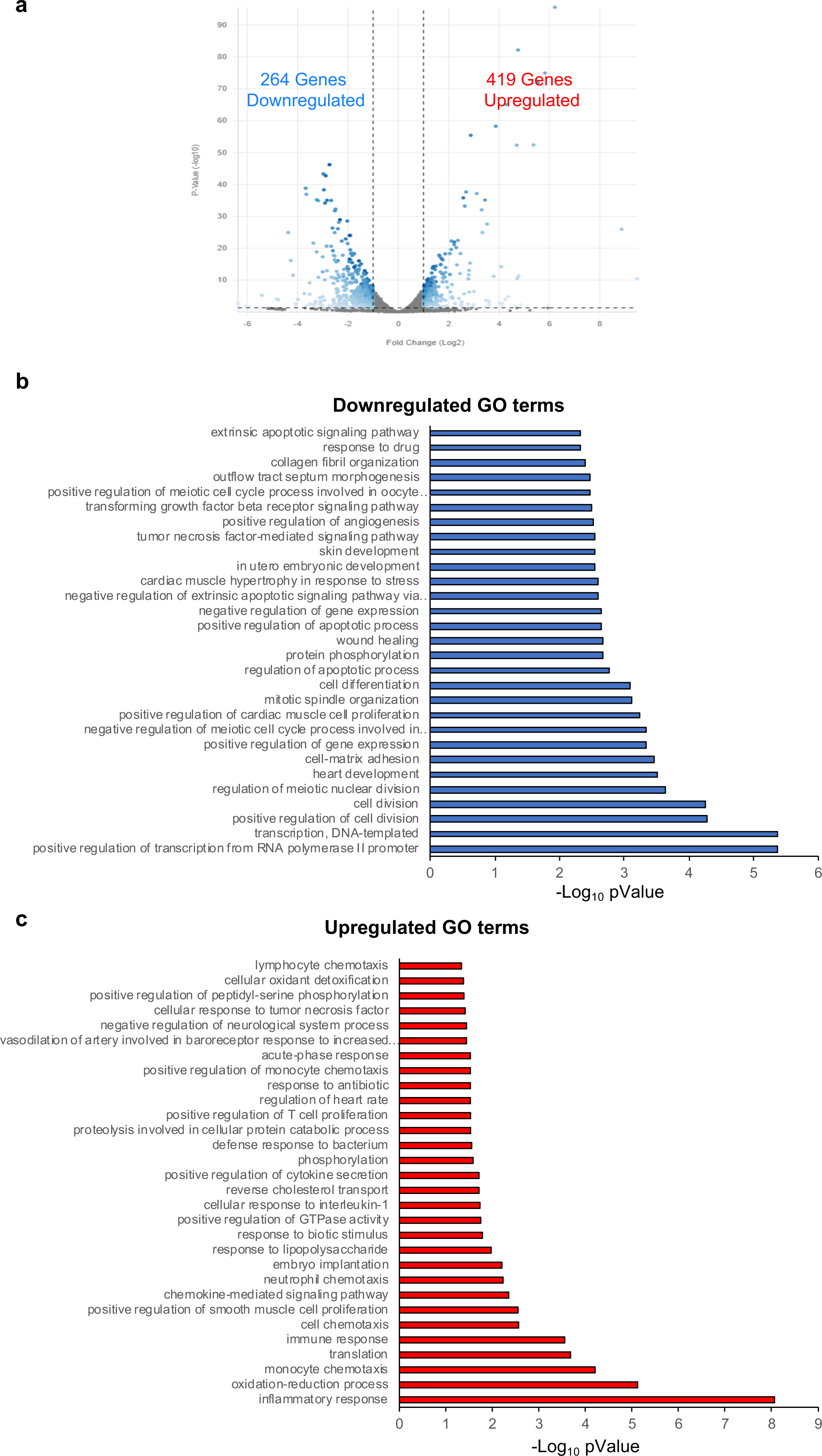
Differential gene expression in slices treated with 100 nM doxorubicin: **(a)** Volcano plot showing significant changes in gene expression in 100nM doxorubicin (Dox) treated tissue. Bar graph shows the GO terms for the significantly downregulated **(b)** or upregulated **(C)** genes from RNA-seq data between control heart slices and heart slices treated with 100nM doxorubicin (n=2 pig hearts).

**Figure 5.**
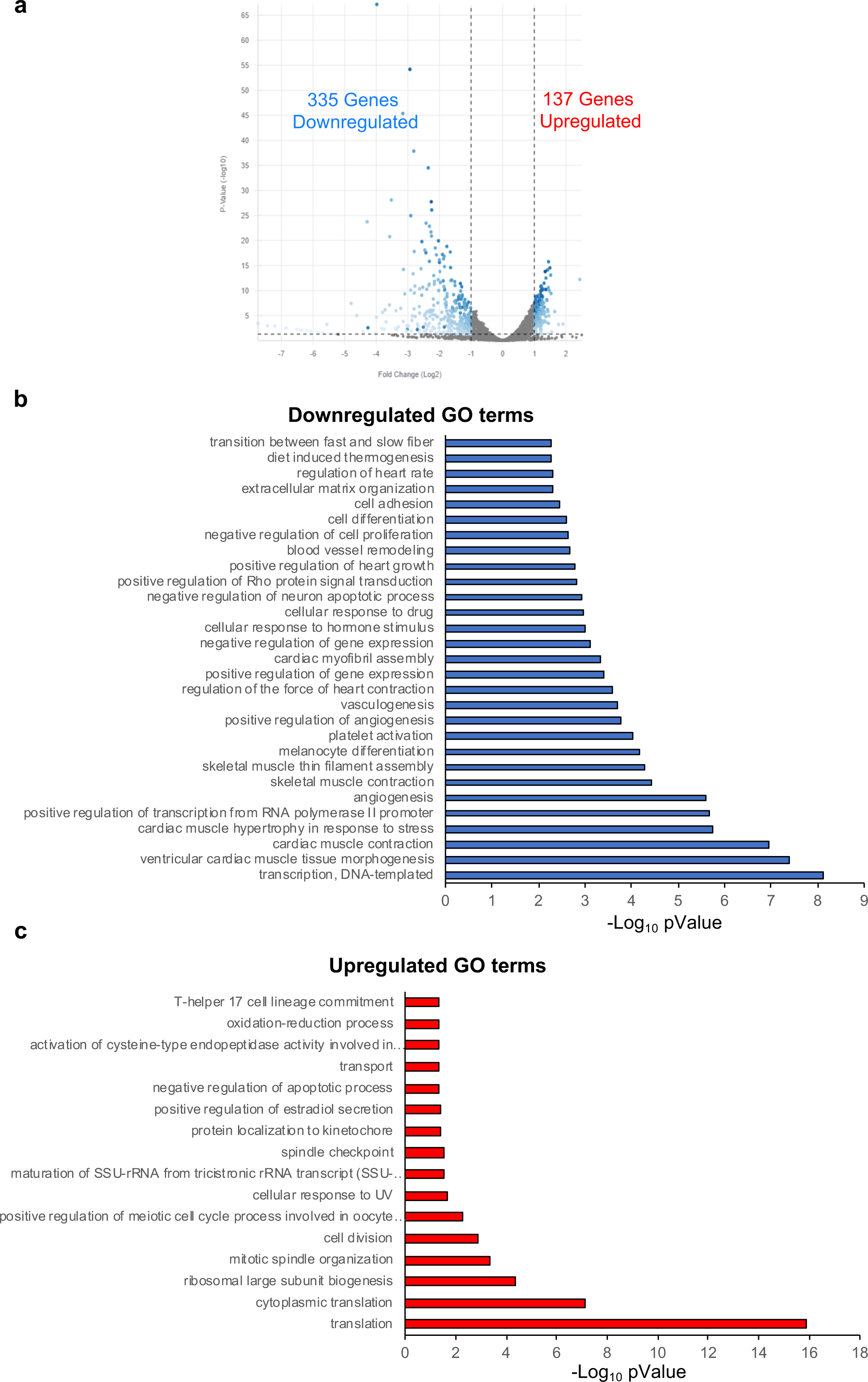
Differential gene expression in slices treated with 1µg trastuzumab: **(a)** Volcano plot demonstrating the genes which are significantly different between control heart slice and slices treated with 1µg trastuzumab. Bar graph shows the GO terms for the significantly downregulated **(b)** or upregulated **(c)** genes from RNA-seq data between control heart slices and heart slices treated with 1µg trastuzumab (n=2 pig hearts).

**Figure 6.**
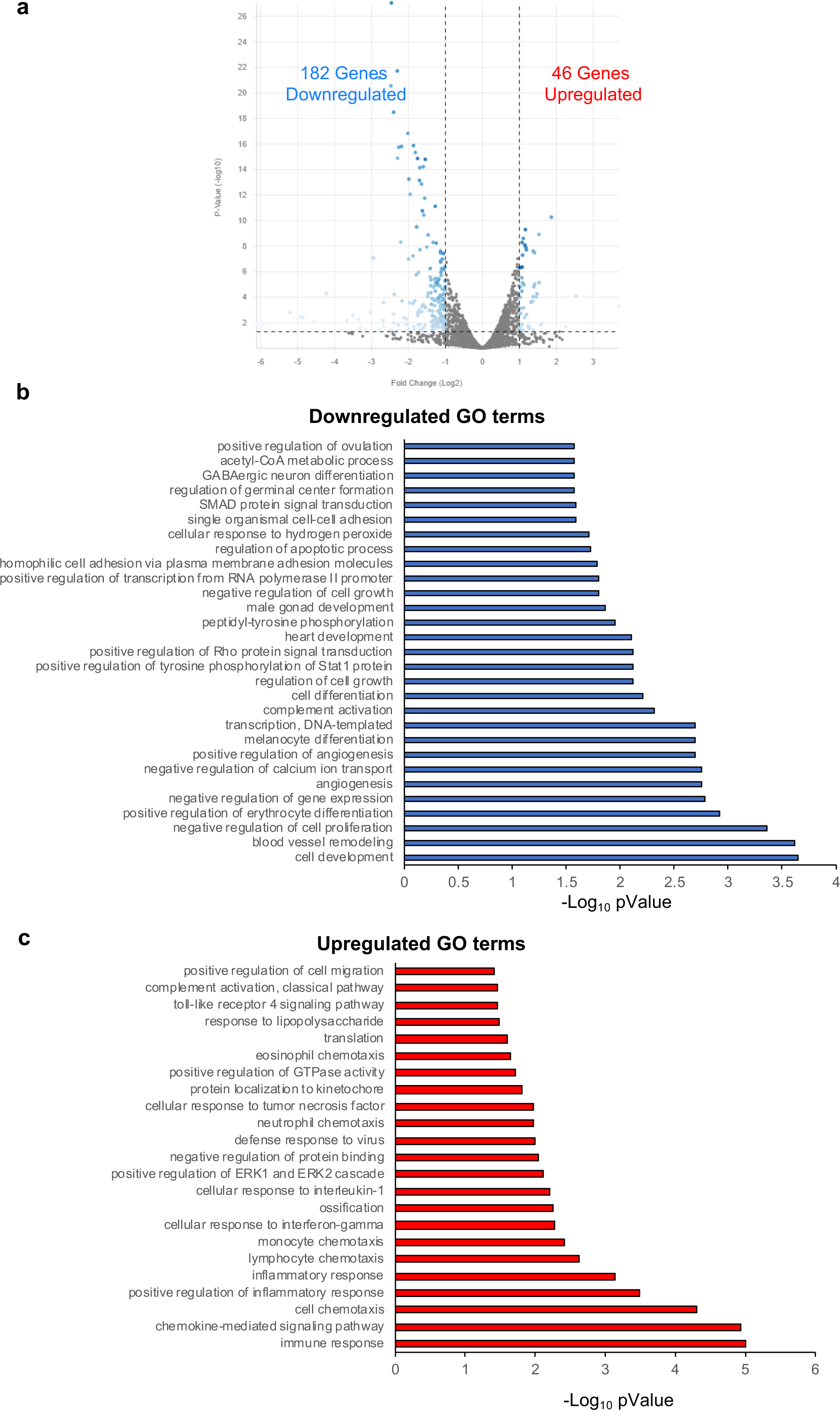
Differential gene expression in slices treated with 100nM sunitinib: **(a)** Volcano plot demonstrating the genes which are significantly different between control heart slice and slices treated with 100nM sunitinib. Bar graph shows the GO terms for the significantly downregulated **(b)** or upregulated **(c)** genes from RNA-seq data between control heart slices and heart slices treated with 100nM sunitinib (n=2 pig hearts).

### Human heart slices can reliably and consistently predict doxorubicin cardiotoxicity

The experiments described above with viability, structure and calcium assays were performed on pig hearts to ensure consistency between the tissues. However, the ultimate goal of the field is to predict cardiotoxicity in human heart tissue, therefore, we used healthy human heart tissue to demonstrate doxorubicin cardiotoxicity on functional and transcriptomics levels. To ensure gender consistency between the human hearts in demonstrating the mechanism of doxorubicin cardiotoxicity, we used 2 female and 4 male hearts. Consistent with the pig heart data, electrophysiological remodeling was also observed in human heart slices. Significant slowing of cardiac conduction velocity in the transverse direction of propagation in response to 50 µM doxorubicin exposure for 24 h was observed in all human hearts used within the experiment^36^ (Figure 7a-b). These results are consistent with reports of acute electrophysiologic alterations in some patients treated with doxorubicin^41^. Furthermore, Cap analysis of gene expression (CAGE) transcriptome analysis revealed 1433 differentially expressed genes in control and 2172 differentially expressed genes at the promoter level in doxorubicin-treated human cardiac slices as illustrated in (Figure 7c). GO analysis revealed that most differentially expressed genes were associated with DNA repair, oxidation-reduction, mitochondria viability and oxidative phosphorylation, which are likely to be related to the damage of the energetic capacity of the cardiomyocytes (Figure 7d). Hierarchical clustering analysis further illustrated different clustering of control and doxorubicin-treated human cardiac organotypic slices (Figure 7e). Importantly, these analyses show the consistency of the transcriptomic response to doxorubicin among different human subjects (Figure 7e).

**Figure 7.**
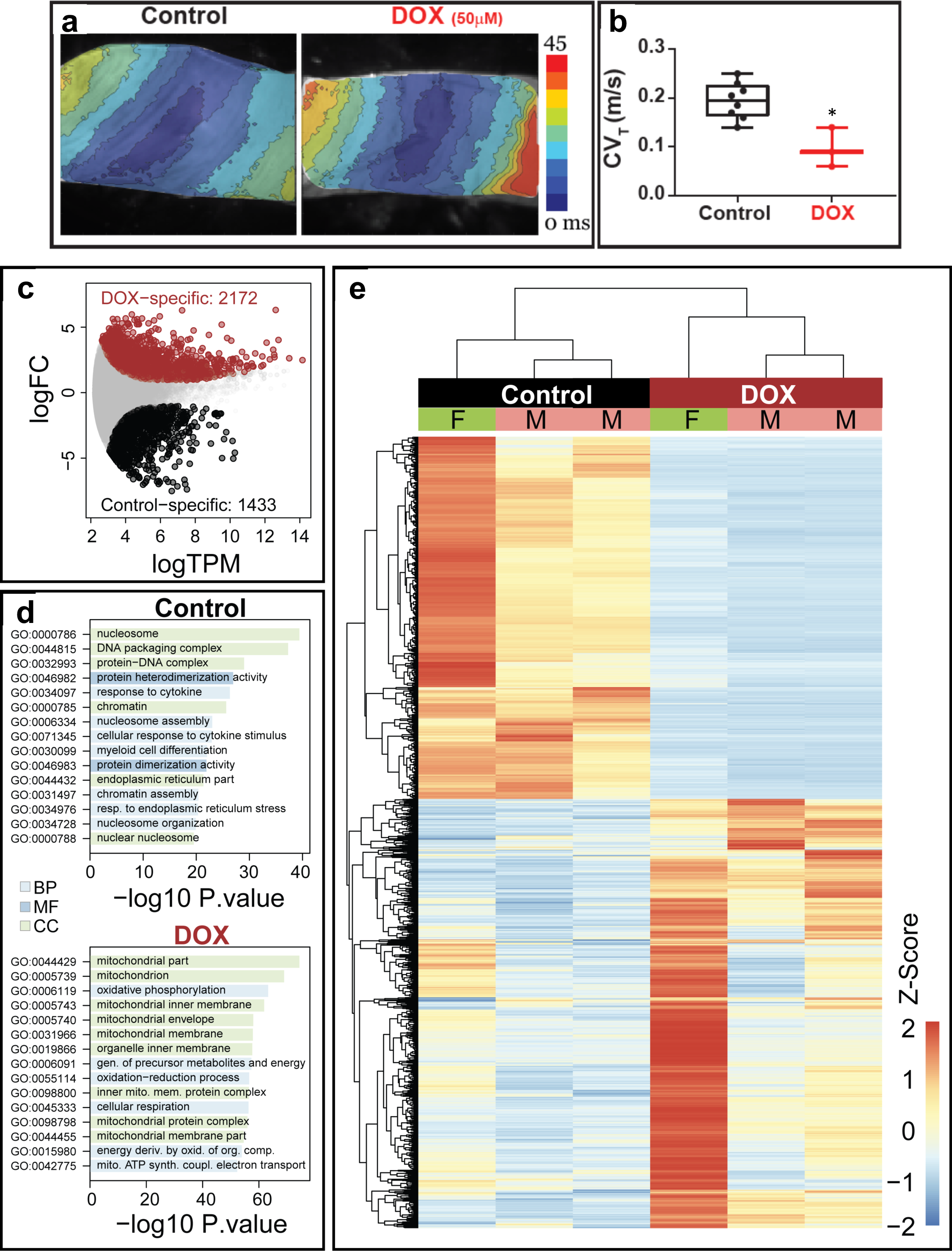
Doxorubicin toxicity in human heart slices: **(a)** Activation maps obtained by optical mapping of human cardiac organotypic slices cultured for ∼24 h with or without doxorubicin (50 µM). Crowding of activation lines in the transverse direction indicate conduction slowing with doxorubicin treatment. **(b)** Average transverse conduction velocity determined from human cardiac slices with and without doxorubicin treatment. Transverse conduction velocity was significantly slower in doxorubicin treated slices (n=7, p<0.05). **(c)** Differentially expressed genes between control and doxorubicin-treated human cardiac organotypic slices following Cap Analysis of Gene Expression (n=3). **(d)** Gene ontology (GO) enrichment analysis of differentially expressed genes in control and doxorubicin-treated slices. BP: biological process, CC: cellular component, MF: molecular function (n=3). **(e)** Heat map demonstrating hierarchical clustering of differentially expressed genes (n=3). F: female, M: male.

## Discussion

Drug-induced cardiotoxicity is a major cause of drug attrition^42^. Therefore, there is a pressing need for predictable preclinical screening strategies for cardiovascular toxicities associated with emerging new drugs prior to clinical trials. The recent consideration of hiPSC-CMs for testing drug toxicity provided a partial solution for some contexts of use but did not solve the overall need in predicting effects in a diversity of functional cardiac modes of failure e.g., effects on endothelial cells, microvasculature or smooth muscle cells^15, 16^. In addition to fetal-like properties, a single cell type does not replicate the complexity of a 3D heart tissue that contains multiple cell types and biological connections. For the first time, we were able to demonstrate the clinical cardiotoxic phenotype of three mechanistically different cardiotoxic drugs using a 3D pig heart slice culture model.

The first drug that we tested was an anthracycline, which is a class of antibiotics discovered over 60 years ago and used to treat many different cancers^43^. Doxorubicin is one of the commonly used anthracyclines for the treatment of lymphoma, leukemia, sarcoma, and breast cancer^43^. The anticancer activity of doxorubicin results from DNA and RNA synthesis disruption^43^, attenuated DNA repair^44^, and generation of free radicals which damage the DNA^45^. Even though doxorubicin is an effective and commonly used anticancer therapy, its use is limited by its cardiotoxicity. Between 5% and 23% of the patients that receive doxorubicin develop diminished exercise capacity and progressive heart failure symptoms^46, 47^. Toxicity mechanisms of doxorubicin dependent on patient-specific properties are still poorly understood, despite being a compound extensively studied for decades; doxorubicin has been found to induce multiple forms of direct cellular injury to cardiomyocytes as a result of free radical production induced by the quinone group^48^. This direct damage to cardiomyocytes has been demonstrated in hiPSC-CMs^28, 30^. Consistent with these reports, our pig and human heart slice models recapitulated the known response to doxorubicin, as our RNAseq data show downregulation of genes responsible for cardiac development and cellular division and upregulation of genes involved in oxidation/reduction and inflammatory response, which is congruent with the known oxidative stress induced by doxorubicin^30^.

The second cardiotoxin is a HER2-targeted agent (Trastuzumab) that targets and inhibit HER2/*neu* receptors (also known as ERBB2). Here we used trastuzumab, the first approved HER2-targeted agent^49^. Trastuzumab is a humanized monoclonal antibody that exerts its anticancer effect through blocking the activation of the HER2(ERBB2)/*neu* receptor, leading to the inhibition of epidermal growth factors/HER2 ligand receptor activity and disrupting the phosphorylation of tyrosine kinases which are critical regulators of the cell cycle^49^. It has been noted that trastuzumab treatment results in asymptomatic cardiac dysfunction and, less often, symptomatic heart failure in some patients^50^. Trastuzumab-induced cardiotoxicity is thought to be driven by disruption of ERBB2/neuregulin signaling in cardiomyocytes, which is critical for normal myocyte growth, survival, and homeostasis^51^. The trastuzumab-induced direct cardiomyocyte damage phenotype and mechanism has been recently modeled in hiPSC-CMs^29,^52. Consistent with these findings, pig heart slices treated with trastuzumab showed direct cardiomyocyte damage, as evidenced by disruption of calcium homeostasis and downregulation of cardiac contractile gene expression.

Sunitinib was the third cardiotoxin tested, a tyrosine kinase and angiogenesis inhibitors (TKIs), which directly inhibits angiogenesis through inhibition of vascular endothelial growth factor (VEGF). This is usually accompanied by cardiovascular toxicity related to increased blood pressure^53^. Other direct cardiac toxicities included cardiac systolic dysfunction, which has been observed in patients treated with sunitinib and sorafenib^54^ and has been attributed to the disruption of the coronary microvascular pericytes leading to hypoxia of cardiomyocytes^55^ and cardiomyopathy. It was not possible to model these indirect effects using hiPSC-CMs, as nanomolar concentrations of sunitinib treatment have no obvious cardiotoxic effects on hiPSC-CMs due to its indirect effect on cardiomyocytes as indicated by our own data (Figure 1) and others^30^. However, in our heart slice system we were able to demonstrate clearly that although sunitinib (as low as 100nM) causes no major damage to the cardiomyocyte structure, it induces disruption of the angiogenic gene program in the heart tissue, which is consistent with its clinical effects.

In conclusion, the results of this study suggest that the heart slice culture system is a promising platform for modeling cardiotoxicity phenotypes and mechanisms. These data suggest that this system will be useful to test acute and subacute cardiotoxicity of new therapeutics.

## Acknowledgments

TMAM is supported by NIH grants R01HL147921 and P30GM127607 and American Heart Association grant 16SDG29950012. The authors also acknowledge NIH grants P30GM127607 (BGH), R01HL130174 (BGH), R01HL147844 (BGH), R01ES028268 (BGH), GM127607 (DJC), P01HL78825 (RB, BGH) and UM1HL113530 (RB), Leducq Foundation RHYTHM grant (IRE), R01HL126802 (IRE), R44HL139248 (IRE) and an American Heart Association Postdoctoral fellowship (19POST34370122) to SAG. We acknowledge the guidance of Ruslan Deviatiiarov from the Institute of Fundamental Medicine and Biology. Kazan Federal University, Russia in the Cap Analysis of Gene expression data analysis.

## Author contributions

J.M.M, M.H.M, Q.O, S.A.G.: collection and analysis of data, manuscript writing, and final approval of manuscript; A.G., R.R.E.A., X-L.T., B.M.A. collection and analysis of data; G.A.G., A.E., B.G.H., J.S., D.J.C., J.M., R.B., A.J.S.R, I.R.E, and T.M.A.M: conception and design, manuscript writing, and final approval of manuscript.

## Competing Interests

TMAM, holds equities in Tenaya Therapeutics. GAG, is consultant for NuPulseCV. The other authors report no conflicts.

## Disclaimer

This article reflects the views of the authors and should not be construed to represent the views or policies of the FDA

## Notes

### Competing Interest Statement

TMAM, holds equities in Tenaya Therapeutics. GAG, is consultant for NuPulseCV. The other authors report no conflicts.
This article reflects the views of the authors and should not be construed to represent the views or policies of the FDA

### Summary of Updates

For a better flaw of the story, we updated the sequence of the figures and edited the manuscript accordingly.

